# Sniffing response to an odor through development in rats: modulatory role of repetition and association with an aversive event

**DOI:** 10.1101/2022.10.17.512510

**Authors:** Julie Boulanger-Bertolus, Emmanuelle Courtiol, Nathalie Buonviso, Anne-Marie Mouly

## Abstract

Sniffing has proven to be a useful behavioral readout for assessing olfactory performance in adult rats. However, little is known about how sniffing response changes through ontogeny. This study thus aimed at characterizing respiratory response to an odor through development in rats using paradigms applicable to both young pups and adults. We first analyzed the sniffing response to the arrival of a novel neutral odor. Then the value of the odor was changed through either its repeated presentation (odor habituation), or its association with a foot-shock resulting in odor fear. In the habituation task, we found that at the three ages, the first presentation of the novel odor induced a clear sniffing response but the peak respiratory frequency was higher in adults than in juveniles and infants. When the odor was presented repeatedly, the sniffing response gradually faded. This habituation of the response took more trials as the animal’s age increases. In the fear conditioning task, the odor induced an increase in respiratory rate that persisted until the end of the session in adults and infants, while it faded rapidly in juveniles. When the odor was explicitly unpaired with the foot-shock, at the three ages the respiratory response to the odor lasted less over the session than in the paired condition. Finally, we observed that shock delivery induced a similar respiratory response at the three ages in paired and unpaired condition. Collectively these data show that sniffing response constitutes a faithful index to assess rat’s olfactory abilities through ontogeny.

## Introduction

When a rat encounters a novel odor in its environment, it initiates an automatic orienting response consisting in an active sampling of that odor achieved via directing its snout toward the odor source and increasing its respiratory frequency resulting in so-called sniffing behavior. As nicely described by Welker (1964) in his seminal study, sniffing in adult rats occurs with a precisely coordinated rhythmic motor sequence involving nose, head, and whisker movements. When they occur together, these movements take place at the same rate and exhibit a fixed temporal relationship to one another. Kurnikova et al (2017) further showed that the onset of each breath initiates a “snapshot” of the orofacial sensory environment. The authors suggest that respiration acts as a master oscillator to phase-lock rhythmic orofacial motor actions. A consequence of the temporal regularity of these signals would be to improve the fidelity of coding of the stimuli.

This stereotypical sniffing behavior undergoes considerable postnatal development. While rapid sniffing is relatively rare and poorly maintained in pups less than 1 week old (Alberts and May, 1980a), by the eighth day after birth, the four sniffing movements are present although not at their maximal amplitude. Between the eighth and tenth postnatal days, exploratory behavior increases and appears to direct the sniffing actions toward target sensory stimuli. Alberts and May (1980) reported that from the second week of life onwards, sniffing becomes a finely-orchestrated pattern of sustained polypnea combined with coordinated movement sequences.

Sniffing has a central role in olfaction since it enhances transport of volatile odorous molecules from the entrance of the nares to the olfactory epithelium. Consequently, sniffing enhances detection and localization of odorants and plays a critical role in odor information processing both in olfactory areas and at higher levels (Buonviso et al. 2006; Mainland and Sobel, 2006; Verhagen et al, 2007; Wesson et al, 2008b). Importantly, odor-induced sniffing proved to be a useful behavioral readout for evaluating olfactory performance in adult unrestrained rats and mice (Macrides et al. 1982; Youngentob et al. 1987; Uchida and Mainen 2003; Kepecs et al. 2007; Wesson et al, 2008a; Courtiol et al, 2014; Lefèvre et al, 2016; Boulanger-Bertolus et al, 2014; Shionoya et al, 2013; Dupin et al, 2020). Odor-induced sniffing has also been used to investigate the ontogeny of olfactory perception in rodents (Alberts and May, 1980b; Boulanger-Bertolus et al, 2014). Indeed, in contrast to classical olfactory tasks that require extensive training and motoric skills that cannot be achieved by pups, odor-induced sniffing provides a reliable index of odor sensitivity in neonates since it involves a spontaneous response to a perceived change in olfactory environment. Using this measure, Alberts and May (1980b) showed that there was a monotonic increase in chemosensitivity as the pups mature from 1 to 17 days of age, with a tendency to a ceiling effect around PN11-13.

Although previous studies have used sniffing behavior to investigate the ontogeny of olfaction during the first weeks of life on one side, and olfactory performances in adult animals on the other side, no study has performed a longitudinal investigation from infancy to childhood with the same olfactory paradigms. The present study was aimed at fulfilling this caveat by comparing the rat’s respiratory response to an odor at different ages of development, using experimental conditions readily applicable to rat pups because they did not require complex movement skills. We first analyzed the sniffing response to the arrival of a novel odor with no behavioral significance. Then the value of the odor was changed through either its repeated presentation leading to odor habituation, or its association with a foot-shock resulting in odor fear. In these two paradigms, respiratory rate was previously shown to be a reliable indicator of the animal’s performance both in pups and in adults (Alberts and May, 1980b; Boulanger-Bertolus et al, 2014; Shionoya et al, 2013). We investigated three ages of development: infant rats (postnatal day PN12-15), juvenile rats (PN 22-24) and adult animals (older than PN75).

## Methods

### Animals

The subjects were male and female Long Evans rats born and bred in the Lyon Neuroscience Research Center (originally from Janvier Labs, France). A different dataset from a subset of these animals has been published in an earlier study (Boulanger-Bertolus et al, 2014). A total of 20 litters were used. Only one female and one male pup per litter per treatment/test condition were used for all experiments and animals from the same litters were used in the different test conditions and ages. Three groups of ages were used: post-natal day 12 to 15 (PN12-15, infants), PN22-24 (juveniles) and older than PN75 (adults). Day of birth was considered PN0. Pups were maintained with their litters up to the end of the experiments, including juvenile pups. Adults were housed by pairs of same sex, at 23°C and maintained under a 12h light-dark cycle (lights on from 6:00 am to 6:00 pm). Food and water were available ad libitum and abundance of wood shavings was supplied for nest building. All experiments were conducted in strict accordance with the European Community Council Directive of November 24, 1984 (84/609/EEC) and the French National Committee (87/848) for care and use of laboratory animals. Care was taken at all stages to minimize stress and discomfort to the animals.

### Training apparatus

The apparatus consisted of a whole body customized plethysmograph (diameter 20 cm, Emka technologies, France) placed in a homemade sound-attenuating cage (L 60 cm, W 60 cm, H 70 cm). The plethysmograph was used to measure respiratory parameters in behaving animals (see Hegoburu et al., 2011 for further description of the plethysmograph). The height of the plethysmograph was adapted to the age of the animal in order to optimize the signal-noise ratio, leading to a height of 30 cm for the adults and 16.5 cm for the infants.

### Odor fear conditioning procedure

Conditioning took place in a sound attenuation chamber with deodorized air constantly flowing through the cage (2 L/min). The odor CS was a 30-s peppermint odor (McCormick Pure Peppermint; 2 L/min; 1:10 peppermint vapor to air) and was controlled with a solenoid valve that diverted the airflow to the peppermint air stream, thus minimizing pressure change. The 1-s mild electric shock was delivered through a grid floor. Adult rats were handled for about 4 days and placed into the conditioning chamber for context habituation. Juveniles received only one day of handling and habituation while infants, for which conditioning to context is not yet developed (Raineki et al., 2010), were not handled to minimize distress from separation from the mother.

Three training conditions were used throughout the experiments (Figure 1): Odor-alone presentations (Odor groups), Odor-shock pairings (Paired groups), Odor-shock unpaired presentations (Unpaired groups). For all groups, animals were allowed a 4min-period of free exploration. Then, in the paired-groups, the CS odor was introduced into the cage for 30s, the last second of which overlapped with the shock. The animals received ten odor-shock trials, with an inter-trial interval of 4min. In the Unpaired groups, the same procedure was carried out except that the shock and the odor were explicitly unpaired using a fixed 180s-interval between the odor onset and the shock arrival. In the Odor groups, the animals received ten 30s-odor alone presentations 4-min apart. In the following, for the three conditions, the term trial will refer to the period starting 30s before odor onset until 210s after odor offset.

**Figure 1:**
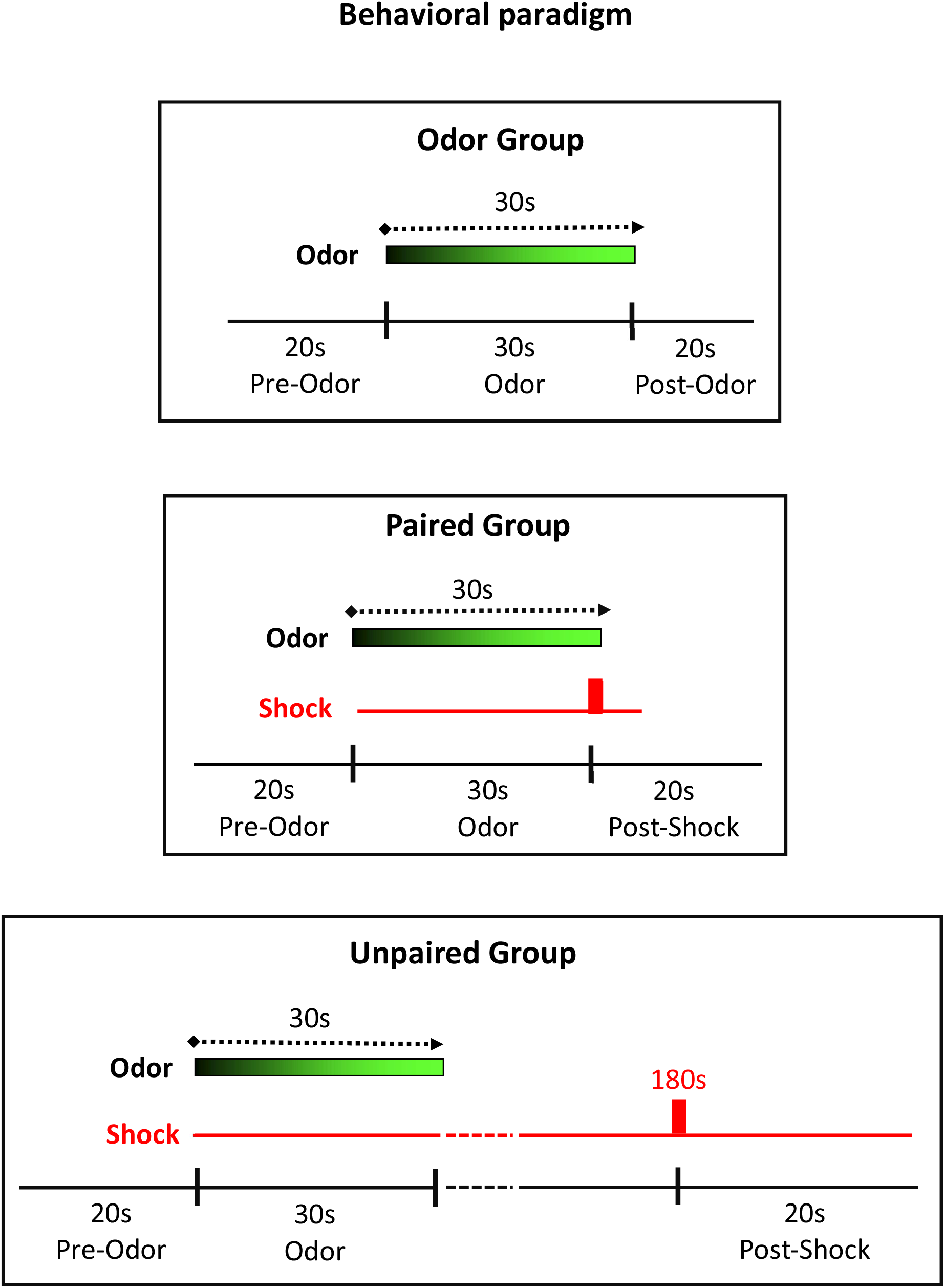
Behavioral paradigms and data analysis periods in the three experimental conditions. Odor group: the animals received 10 presentations of an odor lasting 30s. Respiratory frequency was analyzed 20s before odor onset (Pre-Odor period), 30s during odor (Odor period) and 20s after odor offset (Post-Odor period). Paired group: the animals received 10 Odor-Shock pairings using a 30 s Odor-Shock interval. Respiratory frequency was analyzed 20s before odor onset (Pre-Odor period), 30s during odor (Odor period) and 20s after shock delivery (Post-Shock period). Unpaired group: the animals received 10 presentations of the odor and the shock with a 180s-interval between the two events. Respiratory frequency was analyzed 20s before odor onset (Pre-Odor period), 30s during odor (Odor period) and 20s after shock delivery (Post-Shock period).

### Data analysis

In each experimental group, respiration was monitored throughout the acquisition session. Offline, the respiratory signal was analyzed and momentary respiratory frequency was determined. Instant respiratory frequency was averaged on a second by second basis, leading to 1-s time bins curves. The resulting individual curves were then averaged among animals of the same experimental group. In each experimental group, for each trial, three analysis periods were defined (Figure 1): In the Odor group, 20s-Pre-Odor, 30s-Odor and 20s-Post-Odor; in the Paired and Unpaired groups, 20s-Pre-Odor, 30s-Odor and 20s-Post-Shock. The average respiratory frequency was calculated for each duration period. In each experimental group, the obtained values were compared using three factors ANOVA (Age, Period, Trial) followed by post-hoc pairwise comparisons when allowed by the ANOVA results. For all the statistical comparisons performed, the significance level was set at 0.05.

## Results

### 1 Respiratory response to a neutral odor (Odor Group)

In this experiment, we assessed the effect of a neutral odor presentation on the respiratory frequency. We compared between the three ages the response to the first odor presentation, then to the repeated presentation of this odor, progressively leading to odor habituation.

#### 1.1 First presentation

Figure 2A shows individual examples of raw respiratory signal at the three ages. Figure 2B illustrates the respiratory frequency curve (mean ±SEM) with a 1-sec time bin. At the three ages, there was a clearcut increase in respiratory frequency in response to odor arrival. However, the slopes of the three curves were different, with a peak frequency reaching respectively 10Hz at 9s in adults, 8Hz at 10s in juveniles, and 7Hz at 17s in infants.

**Figure 2:**
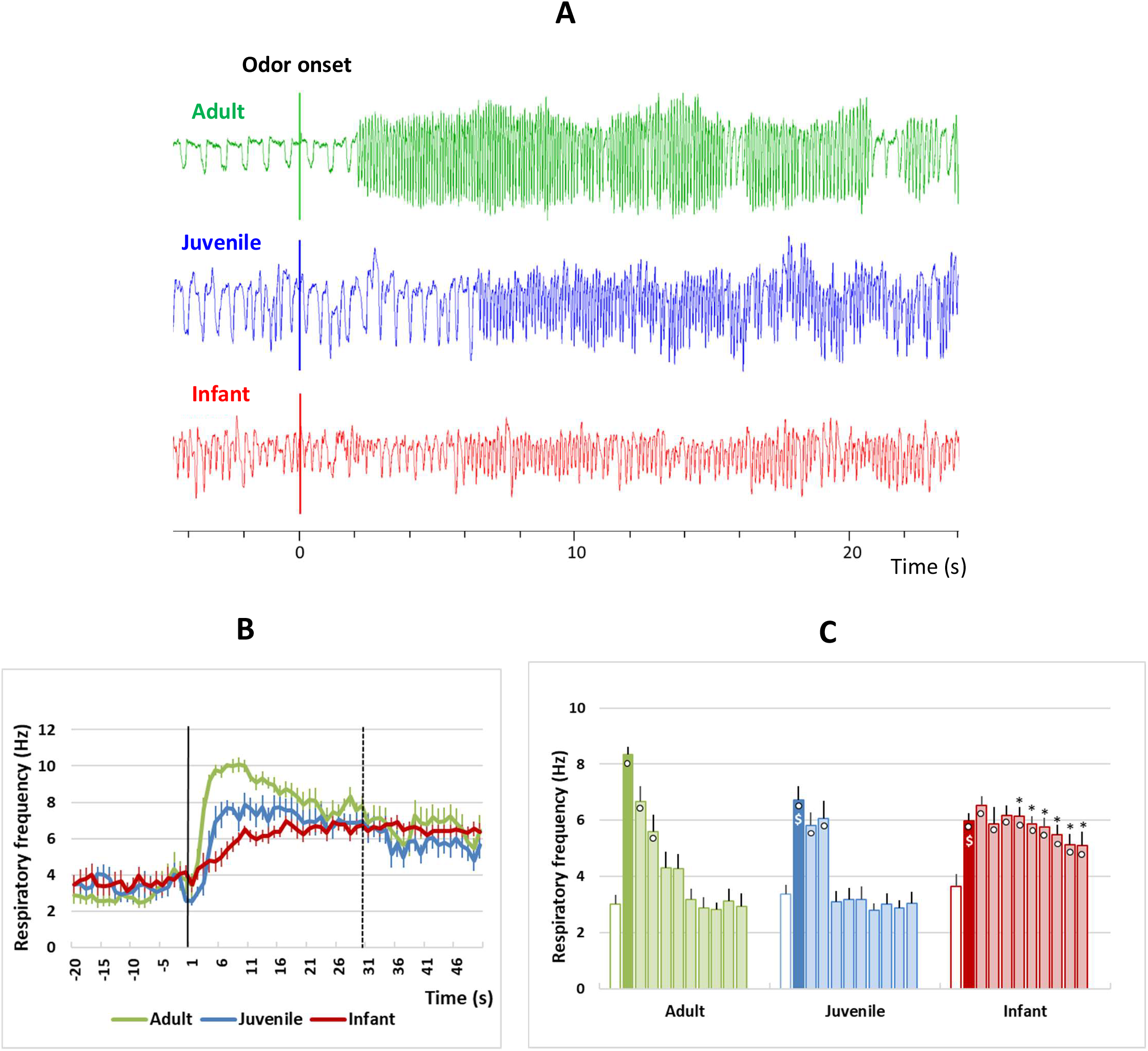
Time course of the respiratory response to the first presentation of an odor across development. A- Individual examples of raw respiratory signals at the three developmental ages in the Odor groups. Following odor delivery, the respiratory frequency increases drastically at the three ages. B- Respiratory frequency time course at the three developmental ages in the Odor group. The time course of respiratory frequency is represented with a 1-s bin precision, from 20 s before odor onset (black vertical line) to 20s after Odor offset (black dotted vertical line). C- Histograms representing the average respiratory frequency during the Pre-Odor (white bars), Odor (plain color bars) and Post-Odor periods (transparent color bars), per 20s time bins until the end of the trial. o: Significant difference with Pre-Odor period; $: Significant difference with Adult at the same period; *: Significant difference with Adult and Juvenile at the same period; p<0.05.

Figure 2C represents the mean respiratory frequency at the three ages during the Pre-Odor, Odor and Post-Odor periods (per 20s time bins) until the end of the trial. A two factors ANOVA revealed a significant effect of Age (F_2,27_=12.26, p<0.001), Period (F_10,270_=35.07, p<0.001) and a significant Period x Age interaction (F_20,270_=6.47, p<0.001). In the three age groups, there is a significant increase in mean respiratory frequency during the Odor period compared to pre-Odor. This increase is greater in adults than in juveniles and infants but is not significantly different between the two younger age groups. During the post-Odor period, in adults and juveniles the respiratory frequency decreases progressively to reach pre-Odor levels from 70s after odor onset onwards in both age groups. In contrast, in infants the respiratory frequency remains significantly higher than pre-odor levels until the end of the trial.

#### 1.2 Repeated presentation

We then looked at the evolution of the mean respiratory frequency before (pre-Odor), during (Odor) and after odor (post-Odor) across the 10 individual trials of the session. Figure 3A illustrates the respiratory frequency curve with a 1-sec time bin at the three ages for the four first trials of the session, the remaining trials being similar to the fourth trial at all ages. Figure 3B represents the mean respiratory frequency for these trials during the pre-Odor, Odor and post-Odor periods. The three factors ANOVA (Age, Period, Trial) carried out on the 10 trials of the session revealed a significant effect of Age, Period, Trial and all the possible interactions (detailed statistics in Table 1). Within group post-hoc comparisons showed that in adults, the mean respiratory frequency during odor was significantly higher than in the pre-Odor period until the third trial. In juvenile animals, the increase during odor lasted until the second trial while in infants it was only observed for the first trial. Thus, the older the animal, the longer it takes for its respiratory response to a neutral odor to habituate.

**Figure 3:**
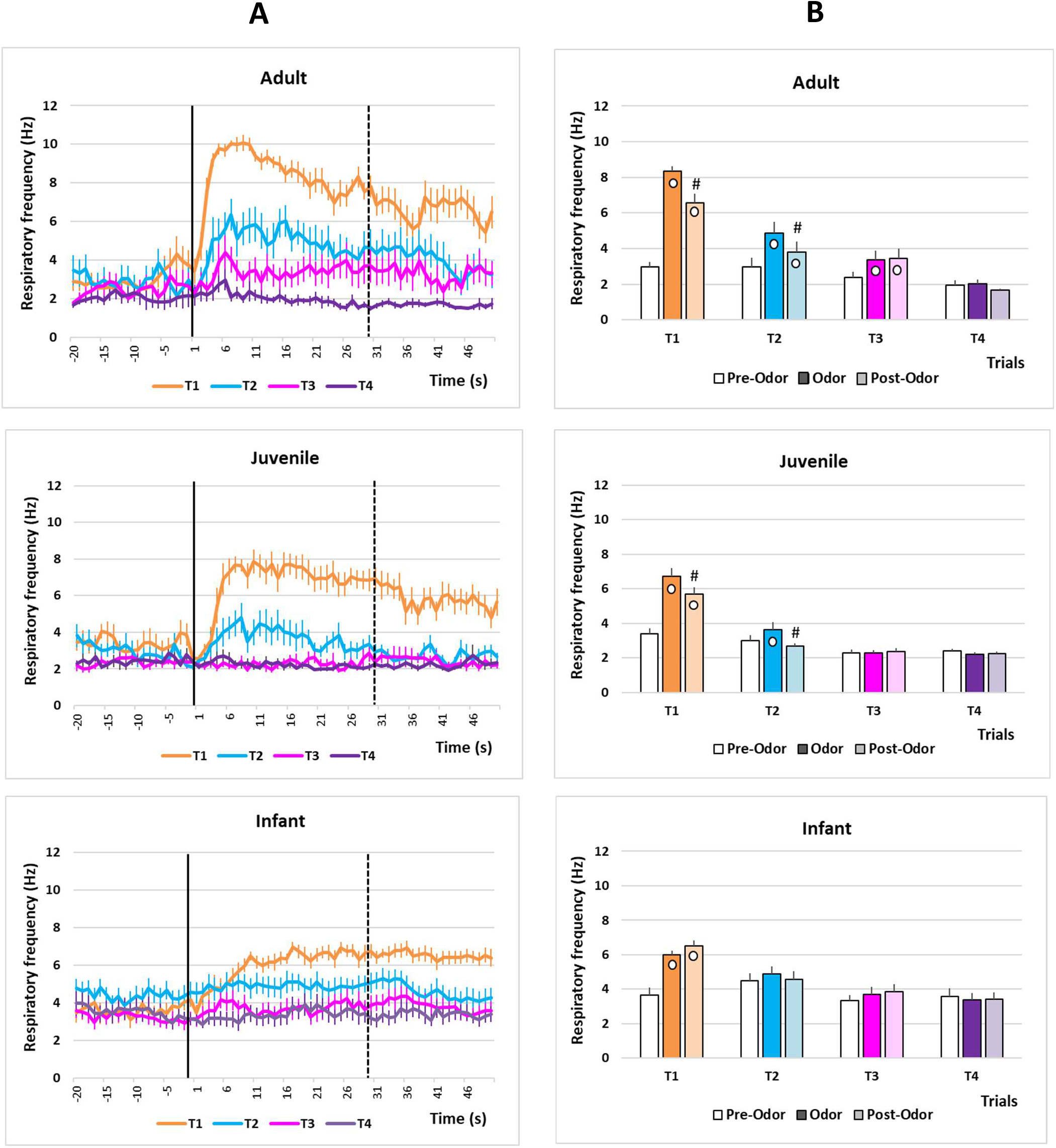
Sniffing response habituation to a neutral odor. A- Respiratory frequency time course for the first four trials (T1 to T4) of the session at the three developmental ages in the Odor groups (from top to bottom: Adult, Juvenile and Infant). The time course of respiratory frequency is represented with a 1-s bin precision, from 20 s before odor onset (black vertical line) to 20s after Odor offset (black dotted vertical line). B- Histograms representing the average respiratory frequency during the Pre-Odor, Odor and Post-Odor periods at the three developmental ages in the Odor groups. o: Significant difference with Pre-Odor period; #: Significant difference with Odor period; p<0.05.

**Table 1:** Statistical results of the three factors ANOVA (Condition, Age, Period, Trial) carried out on the data from the Odor groups presented on Figure 3B. Numbers in red signal significant values of p.

| **Odor group** | **Pre-Odor vs Odor vs Post-Odor** |  |
| --- | --- | --- |
| **Respiration** | **F** | **p** |
| ***Between subjects*** |  |  |
| Age | F(2,27)=4,396 | 0.022 |
| ***Within subjects*** |  |  |
| Period | F(2,54)=82.75 | <0.001 |
| Period*Age | F(4,54)=10.44 | <0.001 |
| ***Within subjects*** |  |  |
| Trial | F(3,81)=68.41 | <0.001 |
| Trial*Age | F(6,81)=3.39 | 0.005 |
| ***Within subjects*** |  |  |
| Period*Trial | F(6,162)=45.28 | <0.001 |
| Period*Trial*Age | F(12,162)=2.43 | 0.006 |

### 2. Response to an odor paired or unpaired with an aversive stimulus

In this experiment, we assessed the respiratory response to an odor in two experimental conditions: either the odor signaled the upcoming arrival of a foot shock (Paired condition) or the odor was not predictive of the footshock delivery (Unpaired condition). In both conditions, we also analyzed the respiratory response to the nociceptive footshock stimulus (i.e. Post-Shock period).

#### 2.1 Paired condition

Figure 4A represents the mean respiratory frequency during the three defined periods across the 10 individual trials of the session, at the different ages in the Paired groups. The three factors ANOVA (Age, Period, Trial) revealed a significant effect of Age, Period and Trial, and all the possible interactions (See statistics Table 2 upper part, for the details).

**Figure 4:**
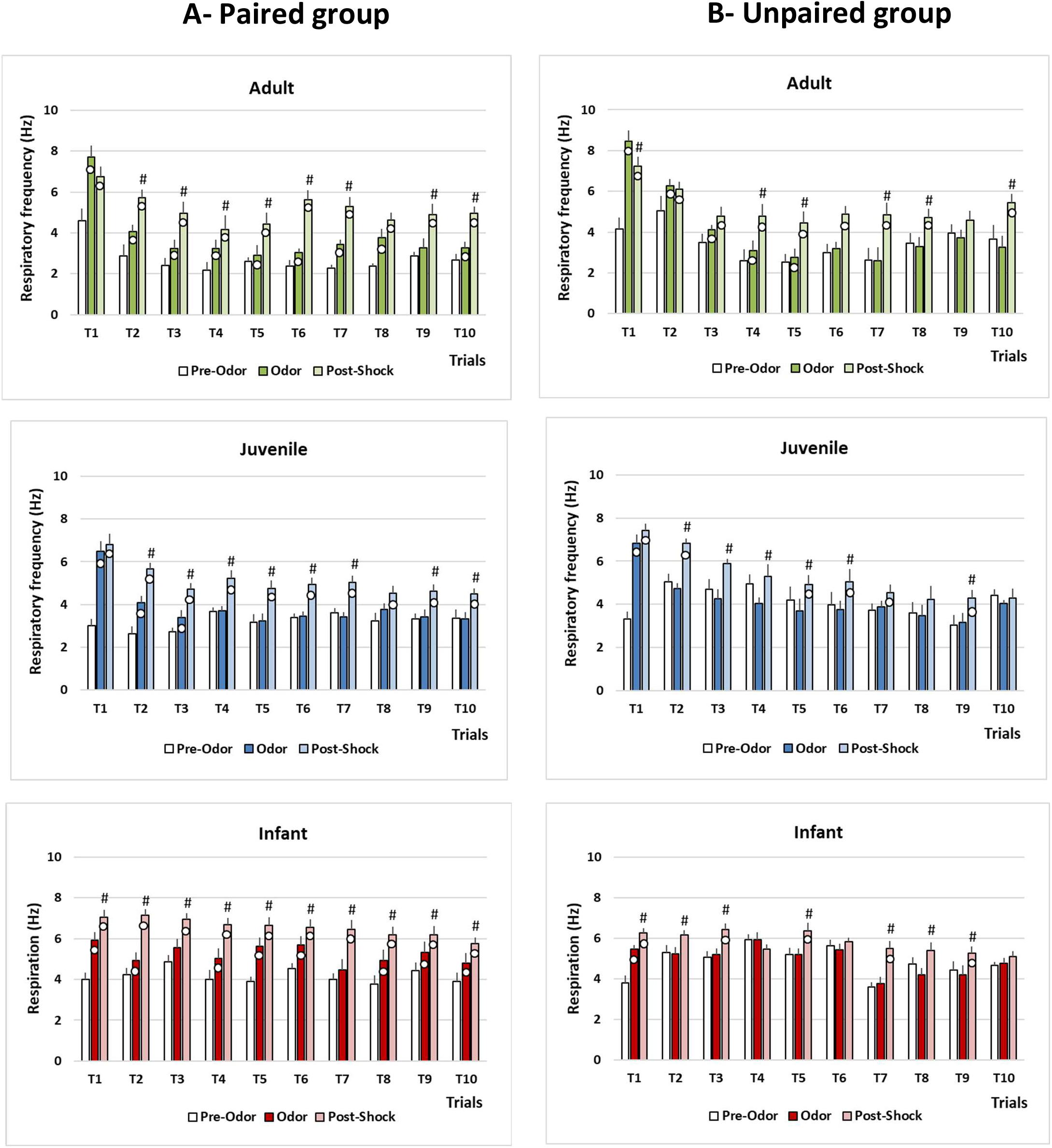
Evolution across trials of sniffing response to an odor in paired and unpaired conditions. A- Histograms representing the average respiratory frequency during the Pre-Odor, Odor and Post-Shock periods for the 10 trials of the session, at the three developmental ages in the Paired group (from top to bottom : Adult, Juvenile and Infant). B- Histograms representing the average respiratory frequency during the Pre-Odor, Odor and Post-Shock periods for the 10 trials of the session, at the three developmental ages in the Unpaired group. o: Significant difference with Pre-Odor period; #: Significant difference with Odor period; p<0.05.

**Table 2:** Statistical results of the three factors ANOVA (Age, Period, Trial) carried out on the data from the Paired (upper part) and Unpaired (lower part) groups presented on Figure 4A and 4B. Numbers in red signal significant values of p.

|  |  |  |
| --- | --- | --- |
| **3 factors ANOVA** |  |  |
| **Paired group** | **Pre-Odor vs Odor vs Post-Shock** |  |
| **Respiration** | **F** | **p** |
| *Between subjects* |  |  |
| Age | F(2,33)=10.35 | <0.001 |
| *Within subjects* |  |  |
| Period | F(2,66)=240.874 | <0.001 |
| Period*Age | F(4,66)=1.786 | 0.142 |
| *Within subjects* |  |  |
| Trial | F(9,297)=18.359 | <0.001 |
| Trial*Age | F(18,297)=4.673 | <0.001 |
| *Within subjects* |  |  |
| Period*Trial | F(18,594)=7.93 | <0.001 |
| Period*Trial*Age | F(36,594)=2.541 | <0.001 |
| **Unpaired group** | **Pre-Odor vs Odor vs Post-Shock** |  |
| **Respiration** | **F** | **p** |
| *Between subjects* |  |  |
| Age | F(2,20)=3.41 | 0.053 |
| *Within subjects* |  |  |
| Period | (2,40)=148.056 | 0.001 |
| Period*Age | F(4,40)=5.352 | 0.002 |
| *Within subjects* |  |  |
| Trial | F(9,180)=16.138 | <0.001 |
| Trial*Age | F(18,180)=4.938 | <0.001 |
| *Within subjects* |  |  |
| Period*Trial | F(18,360)=10.126 | <0.001 |
| Period*Trial*Age | F(18,360)=2.844 | <0.001 |

Within age group’s post-hoc comparisons showed that at the three ages, arrival of the odor induced an increase in the mean respiratory frequency compared to the pre-Odor period. While this increase was maintained for most of the trials of the session in adults and infants, it was only observed until the third trial in juveniles. In regards to the respiratory response to shock arrival, it was similar at the three ages: shock delivery induced a significantly increase in respiratory frequency compared to the pre-Odor and Odor periods for all the trials until the end of the session.

Between ages comparisons carried out on the mean values obtained by averaging all the trials of the session showed that in the three defined periods (pre-Odor, Odor, post-Shock), respiratory frequency was higher in infants than in adults and juveniles (Figure 5A).

**Figure 5:**
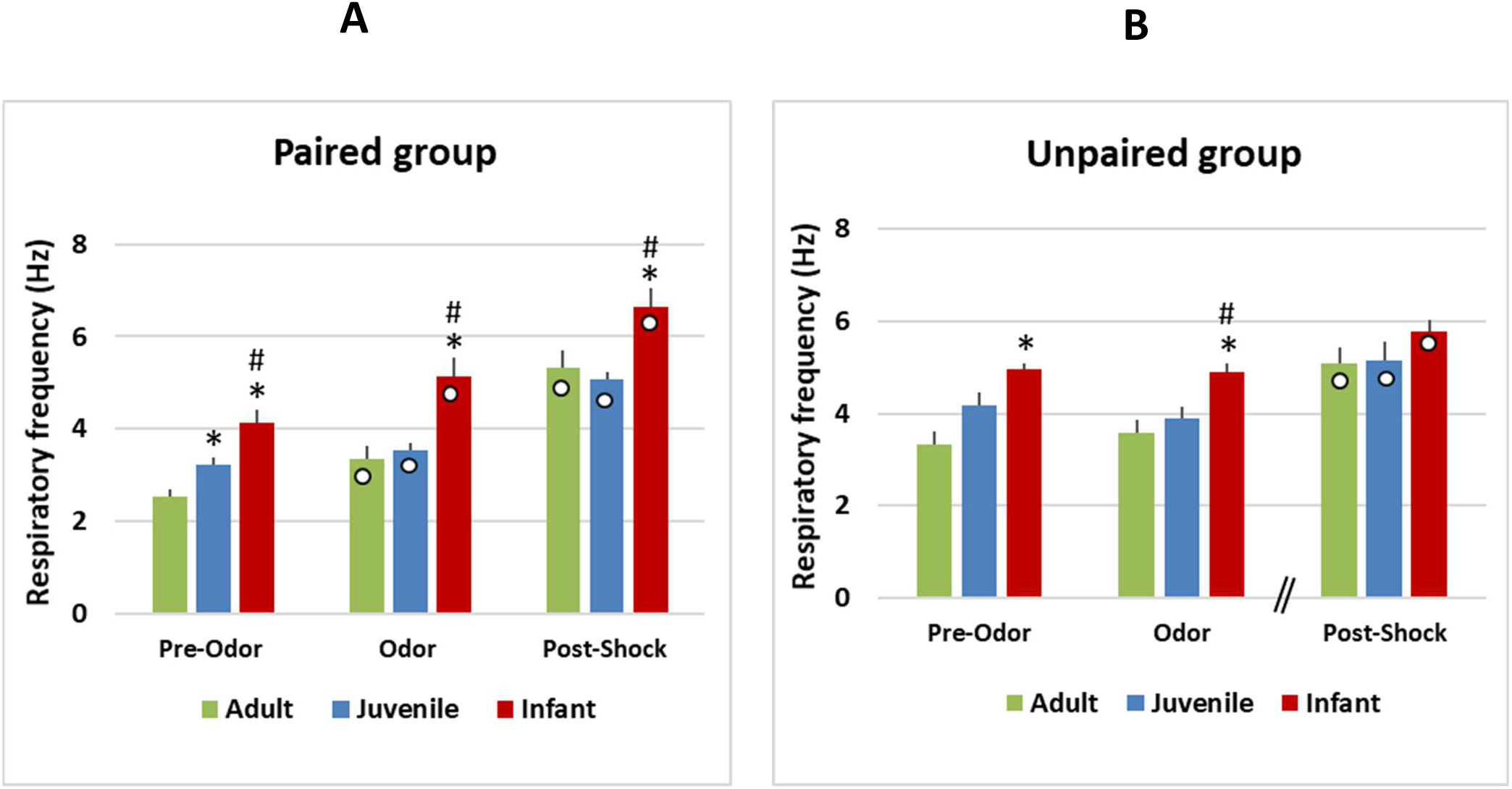
Mean respiratory frequency during paired and unpaired sessions. A- Histograms representing the average respiratory frequency during the Pre-Odor, Odor and Post-Shock periods over the whole session, at the three developmental ages in the Paired groups. B- Histograms representing the average respiratory frequency during the Pre-Odor, Odor and Post-Shock periods over the whole session, at the three developmental ages in the Unpaired groups. *: Significant difference with Adult at the same period; #: Significant difference with Juvenile at the same period; ○ : Significant difference with Pre-Odor period at the same age; p<0.05.

In summary, when an odor is explicitly paired with a foot-shock, arrival of the odor induces a transient increase in respiratory frequency that was observed for each trial until the end of the session in adults and infants, while it vanishes rapidly in juveniles. In contrast, shock delivery induced a similar response at the three ages, consisting in a further increase in respiratory frequency for all the trials.

#### 2.2 Unpaired condition

Figure 4B represents the mean respiratory frequency during the three defined periods, across the 10 trials of the session, at the different ages in the Unpaired groups. The three factors ANOVA revealed a significant effect of Age, Period and Trial, and all the possible interactions (See statistics Table 2 lower part, for the details). Within age group’s post-hoc comparisons showed that in adults, the mean respiratory frequency during odor was significantly higher than in the pre-Odor period until the fifth trial. In contrast, in juveniles and infants the increase during odor was observed for the first trial only. On the other hand, the respiratory response to shock arrival was similar at the three ages: the respiratory frequency after shock delivery was significantly higher than in the pre-Odor and Odor periods for most of the trials of the session.

Between ages’ comparisons carried out on the mean values throughout the session showed that as for Paired animals, in the pre-Odor and Odor periods, respiratory frequency was overall higher in infants than in adults (Figure 5B).

In summary, the respiratory response to the odor lasted longer over the session in Paired than Unpaired animals at all ages. In contrast, the respiratory response to the shock was similar in Paired and Unpaired groups and consisted in a further increase in respiratory frequency for most of the trials of the session.

## Discussion

This study was aimed at characterizing the respiratory response to an odor through development in rats using the same behavioral paradigms. The initial value of the odor was changed through either its repeated presentation leading to odor habituation, or its association with a foot-shock resulting in odor fear. These two paradigms were readily applicable to rat pups because they do not require complex movement skills. In the habituation task, we found that at the three considered ages, the first presentation of the novel odor induced a clear sniffing response. The peak respiratory frequency was higher and occurred earlier after odor onset in adult animals than in juvenile and infant animals. Despite being smaller in amplitude, the sniffing response of infants lasted longer after odor onset than that of adults or juveniles. When the odor was presented repeatedly, the sniffing response gradually faded and then disappeared. This habituation of the response took more trials as the animal’s age increases. In the odor fear conditioning task where the odor signaled the upcoming arrival of a foot-shock, the odor induced an increase in respiratory rate that persisted over the trials until the end of the session in adults and infants, while it faded rapidly in juveniles. When the odor was explicitly unpaired with the footshock, the respiratory response to the odor lasted less long over the session than in the paired condition at all three ages. Finally, we observed that shock delivery induced a similar respiratory response at all three ages in paired and unpaired condition, consisting in a further increase in respiratory frequency in most trials.

Using sniffing response as an index of learning allowed us to detect subtle differences within the conditioning session that the classical measure of freezing would not have unveiled. Indeed, while freezing is a robust and easily quantifiable response, it lacks temporal sensitivity and plasticity. Once induced in response to a foot-shock, freezing often persists throughout the session thus precluding the observation of subtle variations in animal’s fear levels. Respiration in contrast is a more phasic signal than freezing and could give access to transient changes due to odor arrival, otherwise overshadowed by constant freezing (Hegoburu et a, 2011; Shionoya et al, 2013).

### Odor habituation task

At all three ages, there was a sniffing response to the first presentation of the odor suggesting a good perception of the stimulus in our experimental conditions. However, the slopes of the three curves were different, with a peak frequency reaching respectively 10Hz at 9s in adults, 8Hz at 10s in juveniles, and 7Hz at 17s in infants. Thus, the peak respiratory frequency was higher and occurred earlier after odor onset in adults than in juveniles and infants. Importantly, the basal respiratory frequency before odor delivery was not different between ages (Adults: 3Hz; Juveniles: 3.4Hz; infants: 3.6Hz), therefore suggesting the sniffing response to the odor selectively changed through development. Alberts and May (1980a) showed that from the second week of life onwards, sniffing behavior components are well orchestrated. Our data suggest that the vigor of odor-induced sniffing is lower in infants and juveniles, but this does not happen at the detriment of odor perception. Interestingly, while in adults and juveniles the sniffing response induced by the odor returned to basal levels 70s after odor delivery, which corresponds to the end of odor as assessed at the experimenter’s nose, the infant sniffing response persisted until the end of the 4min-trial. This is in accordance with data showing that between 10 and 15 days of age, pups exhibit sustained exploratory behaviour to a novel environment that may persist for several minutes, and they also show hyperreactivity to novel stimuli (Campbell and Spear, 1972; Bolles and Woods, 1964). Thus, in the present study, the persisting sniffing response observed in infants, is rather the result of the intense exploration triggered by the odor than an index of odor sampling.

Upon repeated presentation of the odor, sniffing response progressively declined over the successive trials. Habituation is classically defined as the progressive reduction of a behavioral response elicited by repeated exposure to a novel stimulus not accompanied by any biologically relevant consequence (Leussis and Bolivar, 2006; McNamara et al, 2008). Habituation is a simple form of non-associative learning that underlies animals’ capacity to tone down their response to familiar inconsequential stimuli in favor of novel potentially more relevant events (Thompson and Spencer 1966). Classically in rodents, habituation to an odor is assessed via measuring the investigation time, defined as the amount of time the animal spent within 1-2 cm of the odorant source with its nose aimed toward it. Over successive presentations of the odor, a progressive reduction in investigation time is observed, indicating that the animal remembers its prior experience with the stimulus and no longer investigates it as if it were novel (McNamara et al, 2008). Although easily applicable in adult animals, this measure lacks precision and sensitivity when young animals are tested. In the present study, using exploratory sniffing as a spontaneously expressed response to novel odorants, allowed us to compare habituation through development.

Upon repeated presentation of the odor, sniffing response in adults was no more detectable by the fourth odor presentation. This is in agreement with data from the literature obtained in unrestrained adult rodents using the sniffing response (Wesson et al, 2008; Coronas-Samano et al, 2016; Al Koborssy et al, 2019). At younger ages, the animals’ respiratory response to the odor habituates earlier during the session: at the third trial in juveniles and second trial in infants. This observation suggests that the intense exploratory period triggered in infants by the first odor presentation habituates rapidly.

It could be argued that this rapid habituation was due to the confounding effect of fatigue, particularly in young animals. This might have been disambiguated using a habituation/cross-habituation task. Indeed, in the habituation/cross-habituation test, animals are presented repeatedly with the same odorant (habituation) after which a new odorant is introduced (cross habituation). An animal is considered as perceiving the second odorant as novel when the exploration time is higher than for the prior trial. In our case, the triggering of a sniffing response by the novel odor would demonstrate that the rapid waning of response to the first odor in young animals was not due to fatigue, but to habituation. In contrast, the absence of a sniffing response to the novel odor, would suggest an effect of fatigue. However, another confounding effect in the habituation/cross-habituation test is the possibility that odor discrimination abilities vary through development and an absence of response to novel odor in infants might be explained by a lack of discrimination between the two odors at that age. In the present study, we assume that the rapid habituation observed in infants cannot be accounted for by fatigue because as discussed below, in the Paired condition, the infants’ respiratory response to the odor did not habituate throughout the session. Thus the present data suggest that young animals tend to classify an odor as familiar more rapidly than adults.

### Odor fear conditioning task

Odor fear conditioning has been well characterized during ontogeny and this literature has shown that fear learning emerges in rat pups around PN10 and is caused by the recruitment of the amygdala during the odor–shock conditioning (Sullivan et al. 2000a; Moriceau et al. 2006; Raineki et al. 2009; Boulanger-Bertolus et al, 2016).

The training paradigm used in the present study in the Paired condition has been shown to result in good odor fear memory at the three developmental ages considered (Boulanger-Bertolus et al, 2014). In contrast to what has been observed in Odor animals, adult and infant Paired animals displayed a sniffing response to the odor until the last trial of the session, suggesting that as the trials accumulate, the odor endowed the value of an arousing/alarming signal predicting shock arrival. In that sense, the sniffing response transitioned from an unconditioned response to the new odor first presentation to a conditioned response reflecting the learned emotional value of the odor. In juveniles, the sniffing response to the odor did not persist after the third trial. Importantly, these animals have been shown to have a good memory of the learned odor when tested 24h later (Boulanger-Bertolus et al, 2014). Therefore, the difference in sniffing response to the odor cannot be ascribed to a difference in quality of learning. Interestingly, based on the developmental literature, the peri-weanling period (PN17–23) is the age range at which fear conditioning to the context emerges (Raineki et al, 2010; Sullivan et al., 2000; Brasser and Spear, 1998, 2004). A particularity of contextual learning at that developmental age compared to adulthood, is that it is displayed by paired but not unpaired animals indicating a potentiation of context learning by cue learning (Raineki et al, 2010; Esmoris-Arranz et al., 2008). In our study, contextual learning might have developed in juveniles as trials accumulate and compete with the odor cue in terms of predictive value, leading to the loss of sniffing response to the odor after the third trial. In contrast, in infants that do not yet learn the context and in adults for which context learning occurs preferentially in unpaired condition, the odor cue keeps a strong predictive value and triggers a sniffing response throughout the whole session.

### Difference between paired and unpaired condition

Our data suggest that until the fifth trial, adult unpaired animals react to the odor in the same way as paired animals. In contrast, from Trial 6 onwards, unpaired animals no longer react to the odor leading us to assume that from there, the odor is no more considered as an alarming signal and could rather constitute a safety signal because it is never causally associated with the shock (Rogan et al, 2005). Although not assessed in the present study, we assume that as classically described in the literature, adults unpaired rats have formed a fear to the contextual cues but not to the odor cue.

In juveniles and infants, no sniffing response to the odor was observed after the first trial suggesting the animals clearly dissociated the odor from the shock. In addition, while no study systematically explored context learning in animals younger than 16 days (our infant animals), previous studies have shown that juvenile rats hardly learn context fear in an unpaired paradigm (Raineki et al, 2010). We therefore suggest that unpaired infants and juveniles behave as control odor animals of the same age with a lack of sniffing response to the odor after trial 1.

### Response to Shock delivery

At all three ages in both Paired and Unpaired groups, shock delivery induced an increase in sniffing response that did not habituate across trials. This increased sniffing response is associated with an ensemble of unconditioned defense responses to the footshock. As nicely described by Fanselow (1982), when a rat receives an aversive electric shock, it reacts with vigorous activity characterized by reflexive paw withdrawal, jumping, and squealing. This activity persists for a brief period and then gradually gives way to freezing behavior. The post-shock activity burst constitutes the unconditioned response to aversive event while post-shock freezing is produced by conditioned fear elicited by cues associated with shock (Fanselow, 1980). Here we show that the post-shock activity burst induced an increase in sniffing behavior that was similar at the three ages, suggesting that perception of the nociceptive stimulus is comparable at the different developmental ages tested in the present study (Collier and Bolles 1980; Barr, 1995; Fitzgerald, 2005). In addition, the effect on sniffing is the same for paired and unpaired animals suggesting that signaling the shock by an odor does not modulate the reaction of the animal compared to an unsignaled shock.

## Conclusion

What does the analysis of sniffing behavior bring to the field of olfactory perception assessment, particularly during ontogeny? Sniffing behavior is a spontaneous unlearned response, functional at early ages of development. This measure allowed us to compare olfactory performances through development using the same index, which would not have been possible using for example classical freezing assessment in the odor fear conditioning paradigm since freezing response is immature in infants. Indeed, in adult rats freezing is defined as an immobile crouched posture with the ventral surface elevated above the floor (Blanchard and Blanchard, 1969). However, while freezing can be observed in postweaning animals (Bolles and Woods, 1964), young preweaning rats are not capable of assuming a crouched posture with their limbs extended to support their body trunk due to their immature musculoskeletal system (Takahashi, 1992), thus precluding the use of freezing behavior as a common index through ontogeny.

Despite its above-mentioned advantages, sniffing behavior as assessed through the lens of respiratory frequency, does not replace overt behavior analysis. Indeed, the behavior of the animal cannot be inferred from its respiratory frequency only since for example during freezing, respiratory rate is around 3-4Hz in rats (Hegoburu et al, 2011; Dupin et al, 2019; Boulanger-Bertolus et al, 2014), which is the same frequency range as that of quiet waking state (Girin et al, 2021). In that case, visual observation of the animal’s behavior is necessary to differentiate between the two states. However, it is important to keep in mind that while the present study was centered on respiratory frequency, other parameters of the respiratory cycle like amplitude, volume, peak flowrate or shape, might bring additional information that could help identifying the ongoing behavior (Youngentob, 1987).

Sniffing response is not exclusively observed in response to an odor. Indeed, it is part of the orienting response and it is triggered whenever the animal detects something new and unexpected in its environment. Sniffing response thus constitutes a useful index to assess rat’s perceptual abilities in different sensory modalities in complement to classical behavior markers.

## Acknowledgements

This work was supported by CNRS-PICS program, Partner University Fund Emotion & Time, LIA CNRS-NYU LearnEmoTime, ANR-Memotime, ANR-TDE, COST-Action TIMELY, NIH-DC009910, NIH-MH091451. This work was performed within the framework of the LABEX CORTEX (ANR-11-LABX-0042) of Université de Lyon, within the program “Investissements d’Avenir” (ANR-11-IDEX-0007) operated by the French National Research Agency (ANR). The authors gratefully acknowledge Ounsa Ben-Hellal for taking care of the animals.

